# Intracortical probe arrays with silicon backbone and microelectrodes on thin polyimide wings enable long-term stable recordings in vivo

**DOI:** 10.1101/2021.04.30.442146

**Authors:** Antje Kilias, Yu-Tao Lee, Ulrich P. Froriep, Dominik Moser, Tobias Holzhammer, Ulrich Egert, Weileun Fang, Oliver Paul, Patrick Ruther

## Abstract

**Objective:** Recording and stimulating neuronal activity across different brain regions requires interfacing at multiple sites using dedicated tools while tissue reactions at the recording sites often prevent their successful long-term application. This implies the technological challenge of developing complex probe geometries while keeping the overall footprint minimal, and of selecting materials compatible with neural tissue. While the potential of soft materials in reducing tissue response is uncontested, the implantation of these materials is often limited to reliably target neuronal structures across large brain volumes.

**Approach:** We report on the development of a new multi-electrode array exploiting the advantages of soft and stiff materials by combining 7-μm-thin polyimide wings carrying platinum electrodes with a silicon backbone enabling a safe probe implantation. The probe fabrication applies microsystems technologies in combination with a temporal wafer fixation method for rear side processing, i.e., grinding and deep reactive ion etching, of slender probe shanks and electrode wings. The wing-type neural probes are chronically implanted into the entorhinal-hippocampal formation in the mouse for *in vivo* recordings of freely behaving animals.

**Main results:** Probes comprising the novel wing-type electrodes have been realized and characterized in view of their electrical performance and insertion capability. Chronic electrophysiological *in vivo* recordings of the entorhinal-hippocampal network in the mouse of up to 104 days demonstrated a stable yield of channels containing identifiable multi-unit and single-unit activity outperforming probes with electrodes residing on a Si backbone.

**Significance:** The innovative fabrication process using a process compatible, temporary wafer bonding allowed to realize new Michigan style probe arrays. The wing-type probe design enables a precise probe insertion into brain tissue and long-term stable recordings of unit activity due to the application of a stable backbone and 7-μm-thin probe wings provoking locally a minimal tissue response and protruding from the glial scare of the backbone.

## Introduction

Current neural probe technologies employed in basic research settings outperform clinically approved implants in terms of lateral resolution and the ability to address larger brain volumes (Herbawi et al., 2018; Raducanu et al., 2017; Steinmetz et al., 2021). However, the chronic applicability of these devices is still limited and strongly dependent on implantation strategy and location (Bjornsson et al., 2006; Kozai et al., 2015a) as well as probe tethering to the skull (Biran et al., 2007). The observed degradation in recording yield is mainly caused by the foreign body response of the tissue in the vicinity of the recording sites (Böhm et al., 2019). This progressive inflammation can be seen as a growing glial scar (Turner et al., 1999) that increases the effective distance between neurons and recording electrodes, and acts like an insulator sheet. Since the distance to the neuronal source (Gold et al., 2006; Henze et al., 2000) and electrical transparency of the interjacent tissue (McCreery et al., 2016; Prasad et al., 2012; Salatino et al., 2017) are the two main factors affecting the signal-to-noise ratio of recorded single cell activity and high frequency oscillations, the recording quality typically substantially decreases with implantation time (Polikov et al., 2005).

Recent studies conclude that several factors potentially reduce or even prevent tissue reaction (Ferguson et al., 2019; Jorfi et al., 2015; Kozai et al., 2015b). Next to the overall implant geometry, which should be kept as small as possible (Seymour and Kipke, 2007; Skousen et al., 2011), the selection of materials directly at the probe-tissue-interface seems to be of particular importance with a preference towards soft materials such as polymers that more closely match the mechanical properties of neural tissue (Lecomte et al., 2018; Liu et al., 2015; Luan et al., 2017). However, the usage of soft materials limits the brain volume accessible since large and complex geometries as required for simultaneous electrophysiological recordings across different brain areas request fixed distances between electrode shanks. Furthermore, the insertion of soft probes asks for the application of dedicated insertion devices, i.e., insertion shuttles (Barz et al., 2019; Chung et al., 2019; Felix et al., 2013), or the use of bio-degradable materials temporarily stiffing the flexible probes (Kozai et al., 2014; Tien et al., 2013), increasing the volume of displaced brain tissue at least during probe insertion and chemically altering the local milieu temporarily around the cells of interest. Alternative approaches aimed at minimizing the local foreign body response at the recording site by applying sophisticated probe geometries that enable small probe structures to be deployed sideways from a larger probe shank (Egert et al., 2013; Massey et al., 2019). Additionally, fish-bone-like, highly filigree probe geometries have been mechanically protected during insertion using a bio-dissolvable stiffening material (Khilwani, 2016; Tien et al., 2013). Unfortunately, no long-term recording data, which could have demonstrated an improved recording yield of these advanced devices, are available from these four studies. (Agorelius et al., 2015) implemented flexible probes with protruding recording sites and showed stable neuronal signal quality for up to 3 weeks. Implantation, however, was only possible to superficial, cortical structures and stiffening required a degradable coating that more than quadrupled the size of the implant.

Consequently, we developed probes that combine the advantages of soft materials at the tissue interface while being backed by stiff silicon shanks that permit the access to deep and distant brain regions. We employed polyimide for fabricating what we term electrode wings that were attached to Michigan-style probe geometries consisting of three probe shanks designed for recordings spanning all hippocampal-entorhinal regions of a mouse brain. Impedance spectroscopy confirmed that the electrodes were in the useful range for local field potential (LFP) as well as single/multi-unit recording (SUA/MUA), and probe insertion tests using an agarose gel-based phantom comparable to the brain tissue confirmed that the probes can be implanted and subsequently removed without being damaged. Finally, we implanted the probes in mice and successfully recorded neural activity (both SUA and LFP) in freely moving conditions for up to 104 days post implantation.

## Materials and Methods

### Recording probes

The standard Michigan-style probe comprises three probe shanks (Figure 1(A), blue probe, length 4.0, 4.8, and 3.2 mm, thickness 50 μm) arranged at a pitch of 0.9 and 0.8 mm and a probe base (width × height: 0.58 × 2.0 mm). Circular electrodes with a diameter of 25 μm are integrated at application specific positions optimal for the targeted brain areas, here optimized for parallel recordings scattered across the entorhinal-hippocampal formation of mice. The pitch among the electrodes varies between 300 μm and 1300 μm. They are realized as protruding platinum (Pt) electrodes deposited and patterned on top of the probe passivation layer and interfaced to the probe metallization through circular vias.

**Figure 1:**
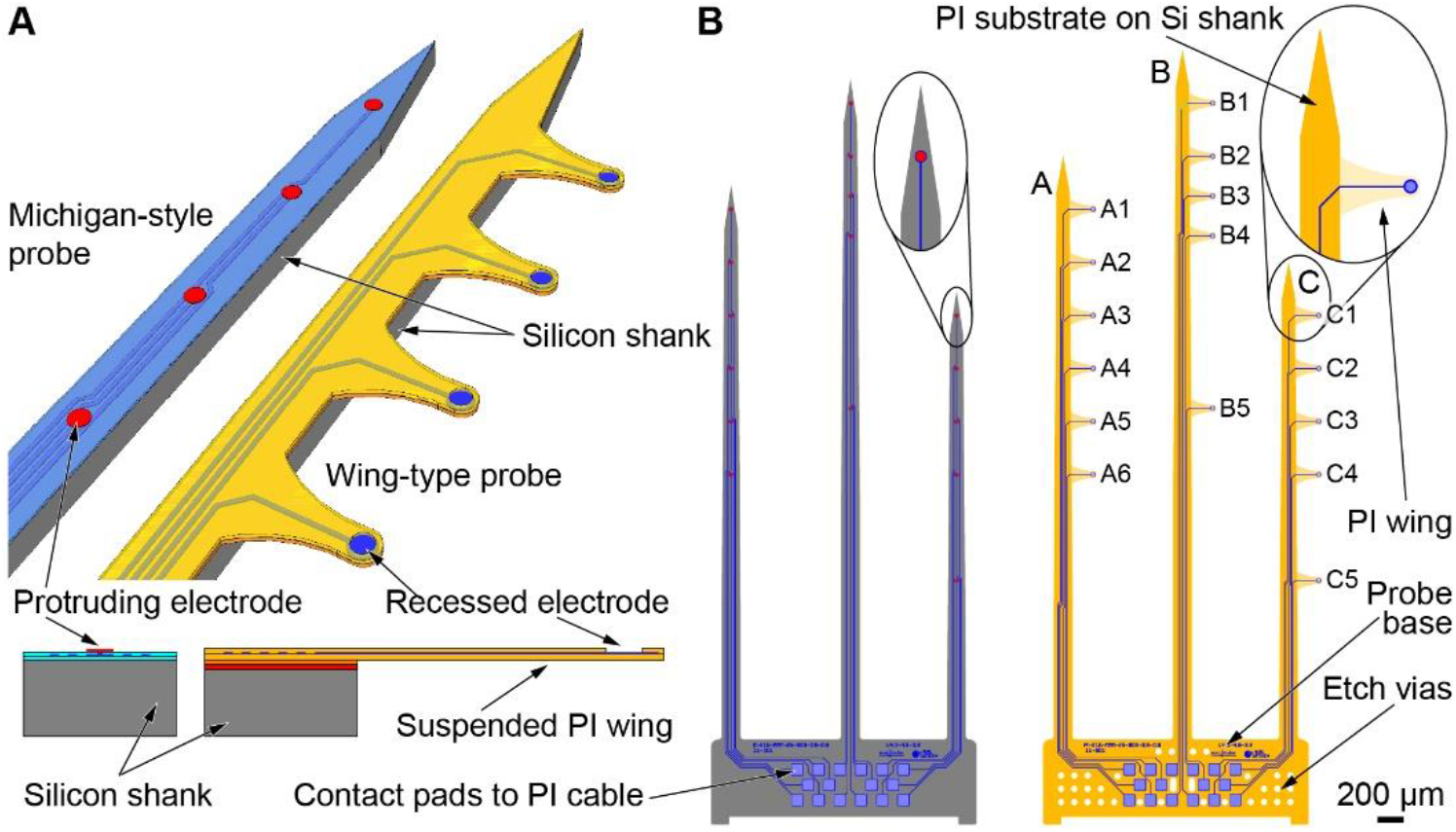
Probe design and comparison to classical Michigan-style probes. (A) 3D schematics and cross-sections comparing the key design features of standard Michigan-style and the wing-type neural probes. The relative electrode positions are identical for both variants. In case of the wing-type probes, electrodes are shifted sideways relative to the supporting silicon shank. (B) Layout of (left) a standard Michigan-style neural probe and (right) wing-type device with electrodes integrated on suspended polyimide (PI) wings. Each probe comprises 16 recording sites distributed along the application specific probe shanks arranged at a pitch of 0.9 mm (shank A-B) and 0.8 mm (shank B-C). The broader probe base comprises respective contact pads interfacing to a highly flexible ribbon cable (not shown) assembled using flip-chip bonding. Etch vias in the probe base of the wing-type probe improve the probe release during processing.

The wing-type probe (Figure 1(A), yellow probe) follow the same basic layout with three probe shanks and a probe base, as well as identical dimensions with regards to electrode position and pitch between shanks and electrodes. The key difference is given by the protruding wing-type, trapezoidal arms with a total length of 200 μm each carrying a single electrode with a diameter of 25 μm. These wings are made of a polyimide (PI) layer stack with a total thickness of 7 μm. As illustrated in Figure 1(B), the wing-type probes apply a Si-based probe comb which is identical to the Michigan-style probe shown in Figure 1(B, left). It serves as a substrate for the PI-based structure comprising the wing-type arms and electrical lines interfacing the electrodes with bond pads on the probe base. In contrast to the Michigan-style probes, electrodes are recessed in the PI sandwich by 3.5 μm (Figure 1(A), cross-section).

Similar to the classical Michigan-style probe applied in chronic settings, the wing-type probe variant is interfaced to the external instrumentation using highly flexible ribbon cables (thickness 10 μm). They are made of two PI layers in-between which a Pt-based metallization (thickness 200 nm) is sandwiched. The respective bond pads of the PI cable have been thickened by gold (Au) electroplating. The cable terminates in a 2×10 strip connector (pitch: 1.27 mm in-line and 2.54 mm between lines).

### Fabrication process

The fabrication process of the silicon-based Michigan style probe is described in detail elsewhere (Herwik et al, 2011). In summary, silicon wafers (diameter 100 mm) with a thickness of 525 μm are coated with a 1.5-μm-thick, stress-compensated layer stack of silicon oxide (SiO_x_) and silicon nitride (Si_x_N_y_) realized by plasma-enhanced chemical vapor deposition (PECVD). Next, the probe metallization made of titanium (Ti)/gold (Au)/Pt/Ti with respective thicknesses of 30/200/100/30 nm is deposited and patterned using sputtering and a lift-off technique based on an image reversal resist (AZ5214E, Microchemicals GmbH, Ulm, Germany). Here, Ti serves as an adhesion promoter to the dielectric PECVD layer while Au is used to reduce the electrical resistance of the metal tracks along the slender probe shanks. The metallization is encapsulated by a second PECVD SiO_x_/Si_x_N_y_ layer stack which is opened at the position of the electrodes and contact pads using reactive ion etching (RIE). The top-most Ti layer is removed by wet etching using hydrofluoric acid (HF, 1%) in order to expose the Pt surface of the contact pads and at the position of the recording sites. Electrodes are realized by the additional sputter deposition and lift-off patterning of a 300-nm-thin Pt layer.

The fabrication technology of the wing-type devices is inspired by the dual sided probe process described elsewhere (Lee et al., 2013). It is summarized in Figure 2 and applies 4-inch, single-side polished Si substrates. The wafers are spin-coated with a 5-μm-thick PI layer (PI2611, HD Microsystems GmbH, Neu-Isenburg, Germany) cured at 350°C (Figure 2(A), PI-1). Next, a 300-nm-thick aluminum (Al-1) layer is sputter deposited after activating the PI surface using an oxygen (O_2_) plasma. It will serve as an etch stop layer during rear side processing. As illustrated in Figure 2(A), the Al-1 layer is patterned by wet chemical etching using a photoresist (PR) mask (AZ1518, Microchemicals). Next, a second 3.5-μm-thick PI layer (PI-2) is spin coated and cured as described before, followed by the deposition and patterning of the probe and electrode metallization (Figure 2(B)). Here, we apply a dual-layer lift-off resist based on LOR (LOA, Microchemicals, thickness 500 nm; technology comparable to (Klein et al., 2018)) and AZ1518 (1.8 μm), followed by surface activation using an O_2_ plasma and physical vapor deposition (evaporation) of Ti (50nm), Pt (230 nm), Ti (50 nm). After spin-coating and curing a third PI layer (PI-3, 3.5 μm, Figure 2(C)), a 10-μm-thick photoresist layer (AZ9620, Microchemicals) is patterned and cured at 115°C for 3 min. The curing process at the elevated temperature creates a curved profile for the electrode opening in the AZ9620 layer. This profile is transferred into the PI-3 layer by a subsequent O_2_-plasma-based RIE process step (Figure 2(D)). This is followed by the sputter deposition of a sacrificial layer (Al, 500 nm (Al-2) and titanium-tungsten (WTi, 200 nm), as illustrated in Figure 2(E). The curved electrode opening ensures that the Al-2 and WTi layers have a good step coverage, which is critical in the final electrochemical release process of the wing-type probes.

**Figure 2:**
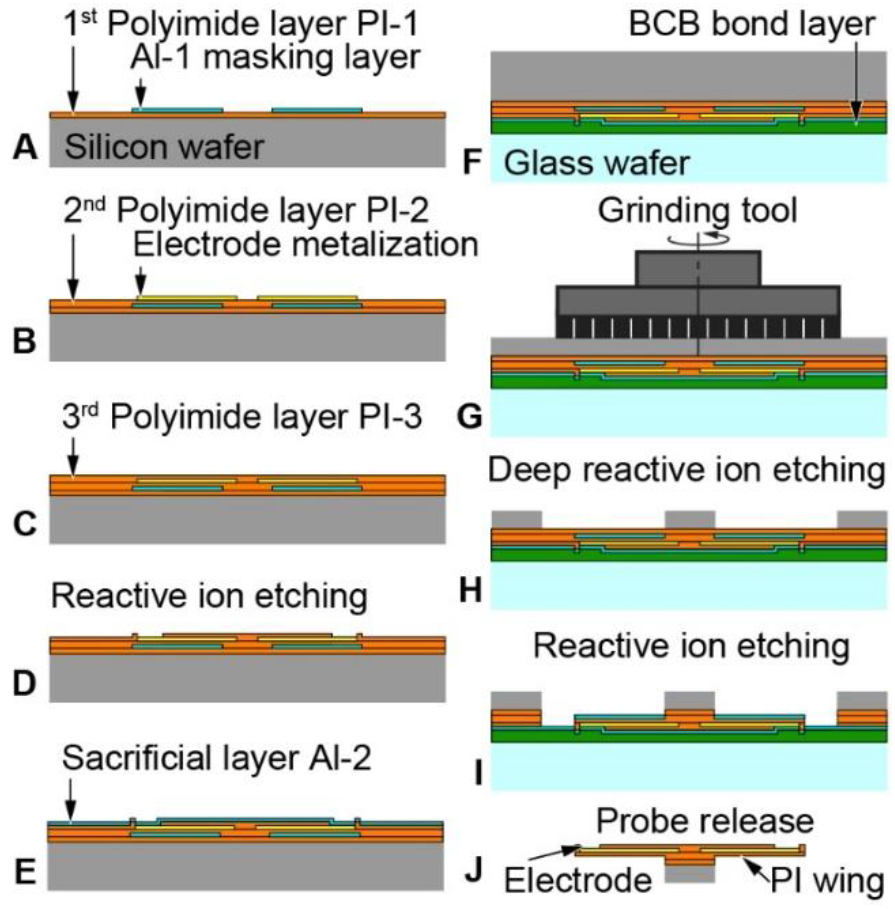
Fabrication of wing-type neural probes: (A) Spin-coating and curing of 1^st^ PI layer (PI-1), and sputter deposition and patterning of Al-based etch stop layer (Al-1, Mask 1); (B) spin-coating and curing of 2^nd^ PI layer (PI-2), and sputter deposition and patterning of metallization for electrodes, tracks and contact pads (Mask 2); (C) spin-coating of 3^rd^ PI layer (PI-3); (D) RIE of PI-3 using a thick AZ9620 layer as masking layer (Mask 3 for electrodes and contact pads); (E) sputter deposition of the Al-based sacrificial layer (Al-2); (F) bonding of the Si wafer to a glass wafer using BCB; (G) wafer grinding to a thickness of 100 μm; (H) DRIE of bulk Si from ground wafer surface using PR as soft mask and PI-1 as etch stop layer (Mask 4); (I) RIE of dielectric and PI layer stacks using Al-1 and Si as masking and etch stop layer; (J) wet etching of sacrificial Al for probe release.

Next, B-staged bisbenzocyclobutene (BCB; Cyclotene 3022-46, Dow Chemical, USA), as initially proposed elsewhere (Niklaus et al., 2001), is spin-coated onto the sacrificial layer to a thickness of 5 μm. The Si wafers are then wafer-bonded to 4-inch Pyrex wafers (thickness 500 μm) using a wafer bonder (MA/BA 6 mask aligner in combination with the SB 6 substrate bonder (both Süss MicroTech AG, Garching, Germany)), as indicated in Figure 2(F). The 200-nm-thick TiW layer deposited onto the Al-2 layer serves as an adhesion promoter to the BCB layer. Similar to the dual sided probe approach described elsewhere (Lee et al, 2013), we used BCB for the intermediate bonding as it is compatible to the DRIE process and offers the optical transparency necessary for the mask alignment of process steps performed on the wafer rear. The applied bonding pressure and temperature were 500 mbar and 250°C, respectively. Similar to earlier work (Herwik et al., 2011; Lee et al., 2013), the Si wafer is then ground from the wafer rear to a nominal thickness of 100 μm. We applied a commercial process provided by DISCO Hi-TEC Europe GmbH, Kirchheim, Germany, as illustrated in Figure 2(G). Using a thick PR (AZ4533, 7 μm) as soft masking layer we apply DRIE to etch through the remaining 100-μm-thick Si substrate and stop on the PI-1 layer. This defines the geometry of the probe shank and base, and removes the bulk Si under the PI wings. This DRIE process step simultaneously introduces etch vias into the probe base (Figure 1(B)), which are subsequently needed for the efficient removal of the sacrificial layer during probe release from the BCB bond layer. Next, RIE is applied to pattern the dielectric layers (PI-1, PI-2 and PI-3) on the Si substrate. The Si shank and the Al-1 layer serve as etching masks to define the shape of the PI wing structures. The etching stops on the Al-2 sacrificial layer which leaves the wafer ready for the probe release.

Finally, the sacrificial Al layer is electrochemically etched in 2M NaCl solution (Klein et al., 2018; Lee et al., 2013; Metz et al., 2005). In order to guarantee electrical contact throughout the entire Al removal process, the WTi layer has been introduction in addition to its adhesion promoting properties. Furthermore, etch vias in the probe base enhance the diffusion-driven transport of the dissolved Al-based sacrificial layer. This electrochemical etch process releases the probes from the PCB layer. Finally, the probes are assembled with a highly flexible PI ribbon cable using flip-chip bonding, as detailed elsewhere (Kisban et al., 2009).

### Impedance spectroscopy

The wing-type electrodes are electrically characterized by measuring their impedance spectra at frequencies between 10^2^ and 10^6^ Hz in Ringer’s solution (Merck, Darmstadt, Germany). A three-electrode setup is used which is comprised of a Pt counter electrode, an Ag/AgCl reference electrode and the working electrode, i.e., the recessed electrode of the wing-type neural probe. The impedance spectra are recorded using an electrochemical impedance analyzer (CompactStat, Ivium Technologies, Eindhoven, The Netherlands) applying a sinusoidal voltage with a peak-to-peak amplitude of 50 mV.

### Probe insertion test

Prior to inserting the wing-type probes into brain tissue, insertion tests were performed using an agarose-gel-based brain phantom. Here, we applied agarose gel prepared from 0.6 wt.-% agarose which was filled into a glass beaker enabling optical inspection of probe insertion using a stereo microscope. The insertion was done by a manually operated linear stage under 90° with respect the flat surface of the agarose gel. The insertion speed was estimated to be 10 mm/min.

### In vivo application

In vivo experiments were performed with adult C57BL/6 mice (8-12 weeks old at implantation, Charles River, Sulzfeld, Germany). All animal procedures were carried out in accordance with the guidelines of the European Community’s Council Directive of September 22, 2010 (2010/63/EU) and were approved by the regional council (Regierungspräsidium Freiburg, Germany). Mice were kept at 22±1 °C in a 12 h light/dark cycle with food and water ad libitum.

Probes were chronically implanted into the hippocampal formation of mice (N = 4; AP = 0.5 mm anterior of the transverse sinuses, ML = 2.7/2.8 mm, DV = 4.5 mm, tilted by 16° towards posterior). Reference electrodes were implanted subcranially above the frontal cortex.

Recordings started after a recovery phase of at least 2 days post implantation. Animals were recorded while exploring a recording cage (34 × 18 cm). Signals acquired by individual electrodes were band pass filtered for 1 Hz - 1 kHz and for 0.4 - 8 kHz to split LFP and MUA. Both signal bands were amplified (LFP: 500×, MUA: 4000×, 2× MPA8I preamps + PGA32, Multichannel Systems, Reutlingen, Germany) and digitized (sampling rated LFP: 2 kHz, MUA: 18.2 kHz; Power 1401 mk2 and expansion ADC16, Spike2 software CED, Cambridge, UK) independently. To assess the performance of the probe quality over time we counted the number of channels showing identifiable SUA or MUA. After the last recording session animals were deeply anesthetized and transcardially perfused with paraformaldehyde (4% solution in 0.1M phosphate buffer). Subsequently the fixed brain tissue was sliced into 40-μm-thick slices. To visualize the astrogliosis around the implant, we performed immunohistochemical stainings against glial fibrillary acidic protein (GFAP, rabbit-anit-GFAP, 1:500, Dako, Hamburg, Germany) counterstained with a Cy3-conjugated secondary antibody goat-anti-rabbit (1:200, Jackson ImmunoResearch Laboratories Inc., West Grove, USA). In addition, we labeled cell nuclei with DAPI (4’,6-diamidino-2-phenylindole; 1:10.000, Roche Diagnostics GmbH, Mannheim, Germany).

Slicing the brain tissue required the probes to be previously removed from the fixed tissue. Explantation of a wing-shaped implant results in post mortal tissue deterioration and extraction of tissue adhering to the probe, both prohibit a detailed analysis of the glial scar or direct comparison to standard probes.

## Results and Discussion

In this study we developed Michigan-style multi-electrode probes on which the recording electrodes were relocated to flexible PI wings. We thereby combined the benefit of a rigid silicon backbone to reach deep brain regions with the advantage of soft materials to minimize the foreign body response and designed a probe optimized for multi-site, chronic *in vivo* recording.

### Processed wing-type neural probes

The fabricated probes implement the concept sketched in Figure 1, i.e., they have the geometry of a 3-shank neural probe with 16 recessed electrodes integrated on 200-μm-long electrode wings protruding sideways from a supporting shank, as shown in Figure 3(A). The probe base comprises etch vias used during probe release via the removal of the Al-based sacrificial layer, as described in Figure 2(I,J). Detailed optical micrographs of wing-type electrodes are shown in Figure 3(B,C) clearly illustrating the interconnecting metal wires. As obvious from Figure 3(C), the electrode wings are straight without observable out-of-plane bending. Furthermore, the hard mask based on Al-1 patterned and applied in the process steps of Figure 2(A) and Figure 2(I), respectively, has been designed a bit larger than the electrode wing itself. The rationale behind this decision was to compensate for a potential lateral misalignment during the photolithography avoiding a gap between the probe shank and the Al-1 masking layer. For this reason, an aluminum strip is left on the silicon shank and sandwiched between polyimide layers PI-1 and PI-2. The inset in Figure 3(C) shows a scanning electron micrograph (SEM) highlighting the sloped sidewall in the PI-3 layer generated in process step (D) according to Figure 2 by applying a thermal reflow process to the PR-based soft masking layer.

**Figure 3:**
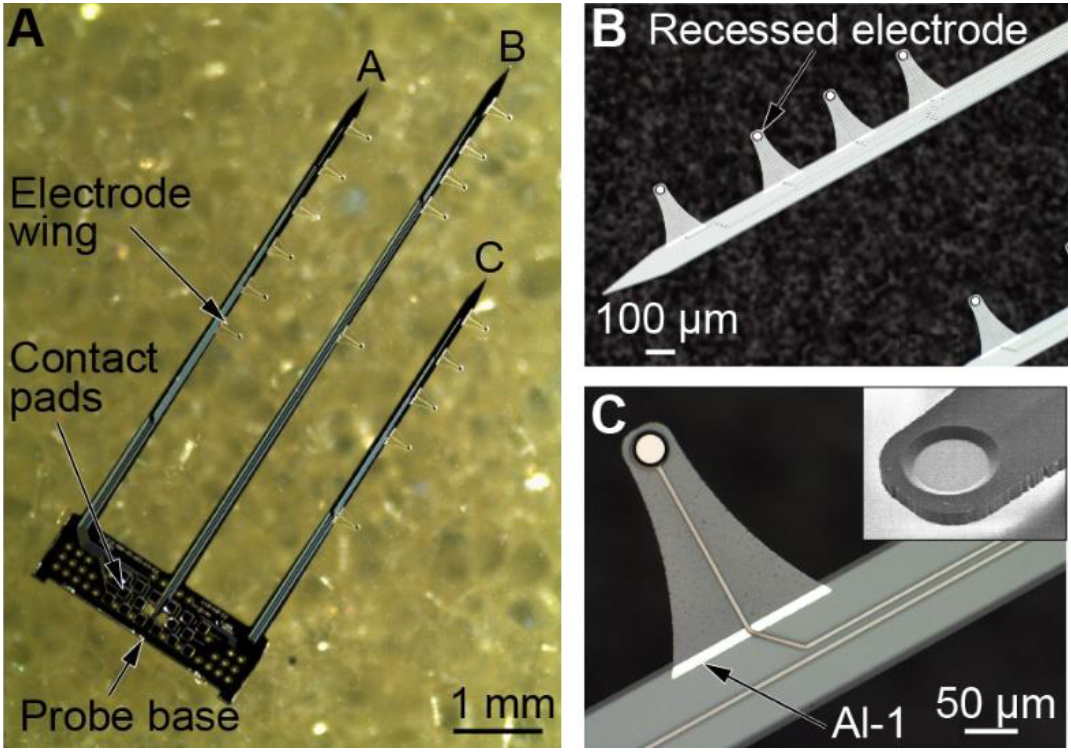
Fabricated wing-type probes: (A) Optical image of a complete probe comb with three probe shanks and 16 electrode wings, and optical micrographs of (B) details of probe shank B with the most distal four wing-type electrodes arranged at a pitch of 400 and 300 μm and (C) detail of a straight wing-type electrode showing a remaining part of the patterned Al-1 layer between layers PI-1 and PI-2 used to define the wing shape in process step of Figure 2(I). The inset in (C) shows a SEM of a recessed electrode with sloped sidewall achieved using the cured AZ9260 PR mask.

### Impedance spectroscopy

Representative impedance spectroscopy data, i.e., absolute value of impedance |*Z*| and phase angle θ, of 15 recessed Pt-based electrodes integrated in protruding PI wings, as shown in Figure 3(C), are given in Figure 4. One electrode of the tested probe with an impedance of 4 MΩ at 1 kHz was rated as defect and omitted from the further analysis. On average, we obtained an absolute impedance value and phase angle at 1 kHz of 325±60 kΩ and 66.8±2.3° (mean ± standard deviation), respectively. The standard deviation of |*Z*| is comparable to Pt electrodes of other Si-based neural probes realized by our group (Seidl et al., 2012). However, with an average value of 325 kΩ at 1 kHz, we observed a reduced impedance which is attributed to a surface roughening of the electrode surface due to the Al-based sacrificial layer, as already described in the case of our dual-sided neural probes (Lee et al., 2013). In future applications, a barrier layer of WTi will be deposited onto the Pt electrodes prior to the sacrificial Al sputtering in order to block the interdiffusion of Al into Pt during the high-temperature PI curing. Nevertheless, with impedances in the 0.3 kΩ range at 1 kHz, the electrodes are in the range useful for neurophysiological application in the high-frequency oscillation range (≤ 250 Hz) as well as for recording SUA/MUA (typically containing frequencies > 400 Hz) (Neto et al., 2018).

**Figure 4:**
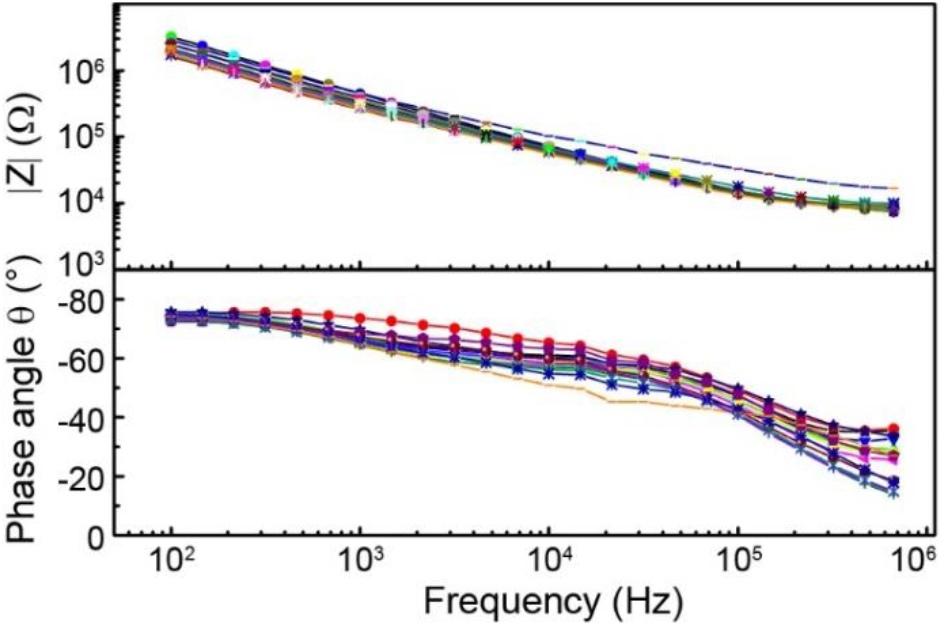
Electrode impedance: Representative impedance spectroscopy data of 15 recessed wing-type electrodes showing (top) absolute impedance |Z| and (bottom) phase angle θ as a function of frequency.

### Insertion test

Figure 5 shows the result of a probe insertion test into an agarose-gel-based brain phantom. The test probe is comprised of two probe shanks each carrying seven wing-type electrodes. The side-view through the transparent agarose gel clearly indicates a safe and bending-free insertion of the 7-μm-thin and 200-μm-long probe wings. Following the insertion test, the wing-type probes were safely removed from the brain phantom without a fracture of the protruding wings. In contrast to the approaches described in (Egert et al., 2013) and (Massey et al., 2019), the symmetric layout of the wings with respect to the implantation and retraction direction is expected to enable the secure probe insertion and removal and to therefore permit recycling of probes for multiple experiments. Admittedly, however, slight tissue damage along the insertion/retraction path due to brain displacement and cutting through the connected brain tissue by the probe shank and electrode wings, respectively, is unavoidable.

**Figure 5:**
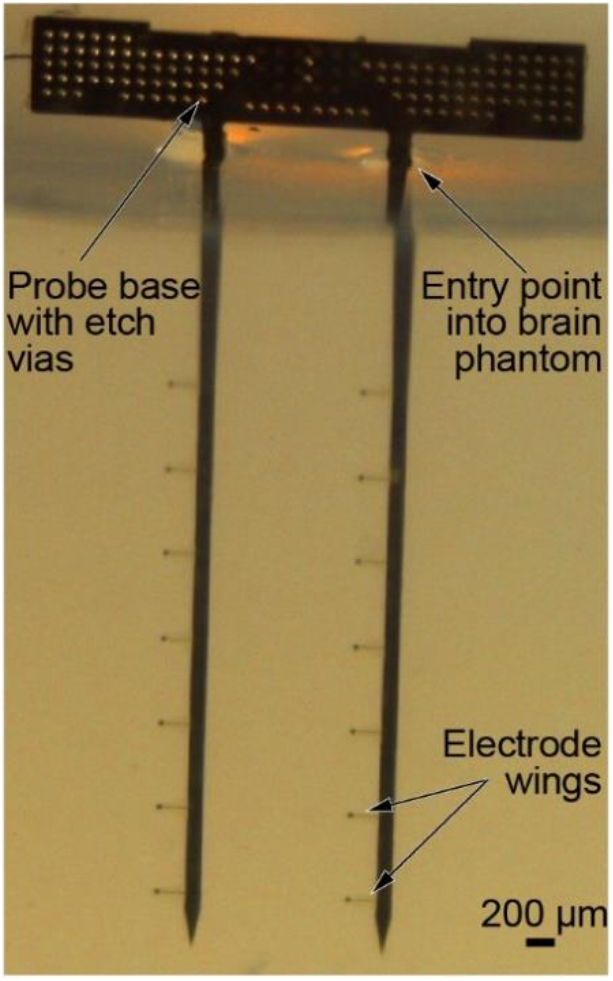
Dummy insertion into brain phantom: Insertion test of a two-shank test probe comprising seven wing-type electrodes per probe shank.

### In vivo application

After confirming the required impedance values and the feasibility of insertion into agarose gel, we tested the applicability of the novel probe design *in vivo* by chronically implanting in total four wing-type probes into mice. A custom-made inserter, as shown in Figure 6(A), allowed us to align the probe on the scull and introduce it, with the intended tilt, deeply into the brain. This enabled recordings of LFP, MUA and SUA at 15 different sites distributed across all hippocampal regions and along the septo-temporal axis of the hippocampal formation (Figure 6(B)). Recordings started 2 days after implantation and were repeated regularly (approximately twice a week) over a time period of up to 104 days. During recordings, mice were allowed to freely explore a recording chamber.

**Figure 6:**
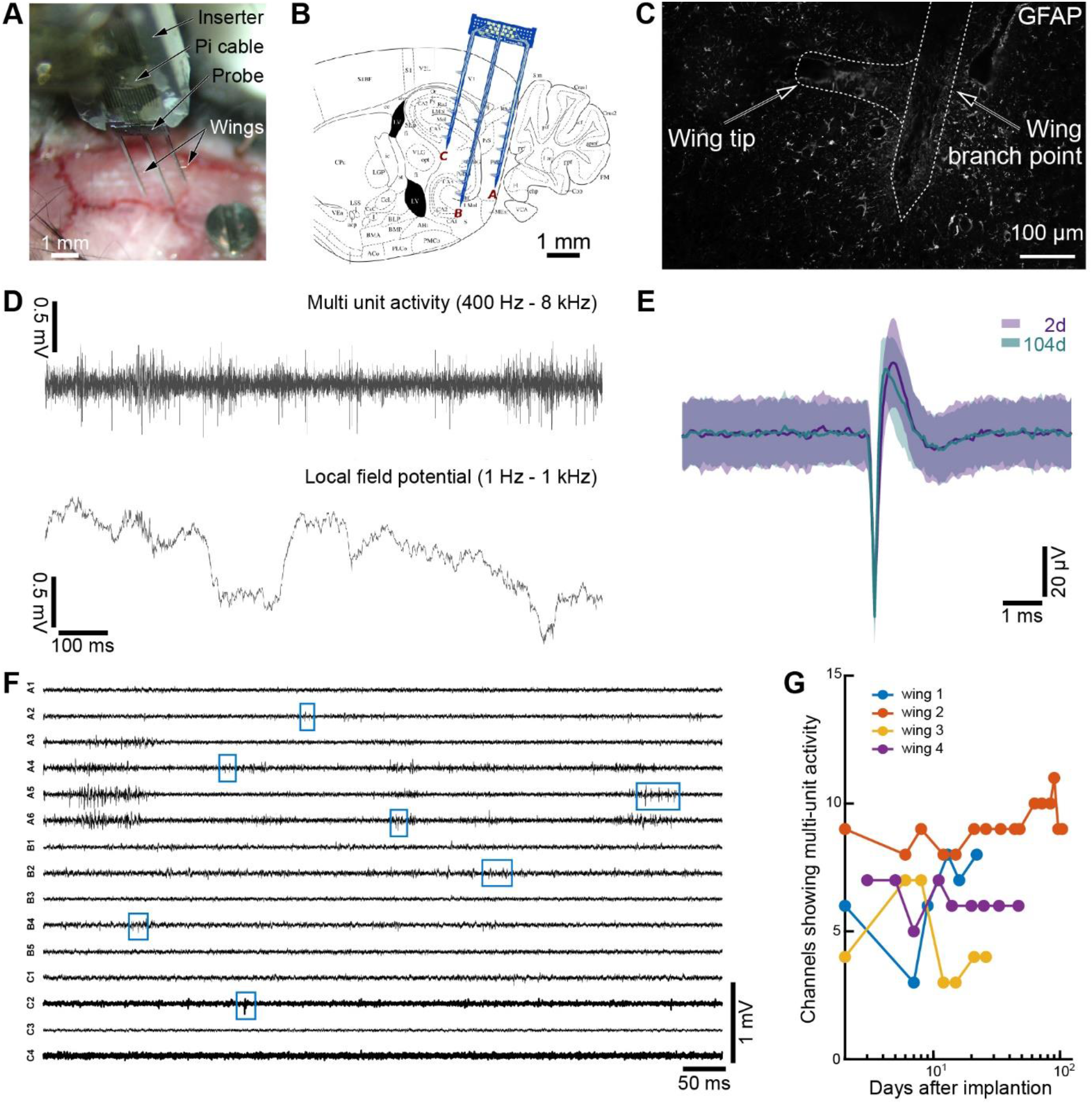
Chronic application of wing-type neural probes implanted in the hippocampal formation of mice. (A) Probe attached temporarily to a custom-made inserter for implantation into (B) different sub-regions of the hippocampal formation. (C) GFAP staining showed a smaller glial response at the tip of the PI wings than close to the main Si backbone. (D) The Pt-based electrodes of reduced absolute impedance allowed for parallel recording of LFP and MUA. (E) Individual unit identified as early as on day 2 after implantation was still identifiable and of comparable amplitude after 104 days of implantation. (F,G) MUA could be seen on a large fraction of channels in parallel (F, blue boxes, wing #2) and this fraction was stable over the implantation duration of each of the four wing-type probes #1 through #4. Wing-type probe #2 was implanted for more than 100 days (recording sites along the probe are identified by a letter and number indicating the shank assignment and electrode position along the shank, respectively, as shown in Figure 1(B)).

To assess the probe-tissue response, we performed GFAP stainings as a marker for the reactive astrogliosis glial reaction and found that at the tip of the PI wings the tissue response was reduced when compared to the response at the silicon backbone (Figure 6(C)). This indicates that the combination of a stiff Si backbone with the soft polymeric substrate material at the recording site is indeed useful for a long-term application in neural tissue. A comparative analysis the reactive astrogliosis would, however, require tissue assess with the probe still in place since explantation retracts or deteriorates the tissue of interest. In particular in high-quality recordings, when neuronal tissue presumably tightly adheres to the recording site, explantation would likely affect the glial scar.

Previous studies have shown that wings with a length of 200 μm should be sufficiently long to protrude from encapsulation of the main Si backbone (Grand et al., 2010) but it would require tissue clearing techniques (Ueda et al., 2020) or 3D-reconstructions of X-ray tomography imaging (Böhm et al., 2019) to verify this.

In terms of neural activities, we observed both LFP and MUA on the same channel using respective filter bands (1 Hz - kHz for LFP, 0.4 – 8 kHz for MUA, Figure 6(D)). The wing-type electrodes even allowed to record from a single unit until day 104 after implantation, as illustrated with the spike sorted data in Figure 6(E). Across all channels we assessed the occurrence of MUA (Figure 6(F)) and used the number of channels that displayed MUA over time as a measure for the long-term stability of the wing-type probes. The fraction of active MUA channels remained very stable over the time assessed in all 4 probes (Figure 6(G)) further supporting the benefits of these novel tools for chronic recording. On average signals from 47.7 ± 14.1% channels contained identifiable MUA/SUA; this yield outperforms probes with electrodes residing on the Si backbone and implementing comparable geometries and electrode materials (Ulyanova et al., 2019).

In the future, in order to adapt the impedance of the probes to the requirements of different applications, surface coatings such as PEDOT for impedance adjustment (Castagnola et al., 2015) or conductive hydrogels or polymers for improved biocompatibility (Aregueta-Robles et al., 2014) could be employed.

## Conclusions

Novel Michigan-style probe arrays comprising polymeric wing-type electrodes that protrude sideways from a stiff, Si-based backbone have been presented. Their fabrication process applies standard microsystems technologies combined with wafer grinding and the thermally stable, temporary bonding of silicon wafers to optically transparent handle wafers. This enables the precise rear side processing of the Si-based support structure of the electrode wings made of PI with a thickness of only 7 μm. As demonstrated with the dual-sided (Lee et al., 2013) and ultra-thin neural probes (Herwik et al., 2011), the Si backbone thickness can be reduced to 50 μm and below, which is expected to further reduce tissue response commonly hindering long-term probe application. Probe insertion into and retraction from an agarose-gel-based brain phantom and mouse brain was possible without damaging the protruding electrode wings. This opens the possibility for probe recycling. However, protruding wings increase the local cross-section of the probe; therefore, tissue damage during implantation caused by cutting through the targeted brain area with the protruding electrode wings is unavoidable. Nonetheless, long-term stable chronic recording from all hippocampal-entorhinal regions of the mouse brain were possible for up to 104 days post implantation with about 50% of all electrodes showing MUA at all times. This high yield supports the notion that the ultra-thin wings effectively cause less tissue damage locally around the recording sites than conventional probe designs.

## Acknowledgement

This work was supported by the National Science Council, Taiwan (NSC100-2627-E-007-002), by the German Federal Ministry of Education and Research (BMBF, FKZ 01GQ0420 and 01GQ0830), and by the Deutsche Forschungsgemeinschaft (SFB TR3 and SFB 780).

## Author’s contributions

YL and PR designed the probes, YL and DM MEMS-fabricated the probes, TH developed the probe assembly to PI-based ribbon cables, AK and UPF performed in vivo tests and data analysis, UE, OP and WF provided scientific support, AK, UPF and PR designed the technical and in vivo study and wrote the initial manuscript, all authors reviewed the manuscript.

